# Reduced Pathogenicity of the SARS-CoV-2 Omicron Variant in Hamsters

**DOI:** 10.1101/2022.01.02.474743

**Authors:** Katherine McMahan, Victoria Giffin, Lisa H. Tostanoski, Benjamin Chung, Mazuba Siamatu, Mehul S. Suthar, Peter Halfmann, Yoshihiro Kawaoka, Cesar Piedra-Mora, Amanda J. Martinot, Swagata Kar, Hanne Andersen, Mark G. Lewis, Dan H. Barouch

## Abstract

The SARS-CoV-2 Omicron (B.1.1.529) variant has proven highly transmissible and has outcompeted the Delta variant in many regions of the world^1^. Early reports have also suggested that Omicron may result in less severe clinical disease in humans. Here we show that Omicron is less pathogenic than prior SARS-CoV-2 variants in Syrian golden hamsters. Infection of hamsters with the SARS-CoV-2 WA1/2020, Alpha, Beta, or Delta strains led to 4-10% weight loss by day 4 and 10-17% weight loss by day 6, as expected^2,3^. In contrast, infection of hamsters with two different Omicron challenge stocks did not result in any detectable weight loss, even at high challenge doses. Omicron infection still led to substantial viral replication in both the upper and lower respiratory tracts and pulmonary pathology, but with a trend towards higher viral loads in nasal turbinates and lower viral loads in lung parenchyma compared with WA1/2020 infection. These data suggest that the SARS-CoV-2 Omicron variant may result in more robust upper respiratory tract infection but less severe lower respiratory tract clinical disease compared with prior SARS-CoV-2 variants.

---

The highly mutated SARS-CoV-2 Omicron variant has led to rapid global spread, including in individuals who have been fully vaccinated^1^. Although case numbers in many countries are currently at unprecedented levels, early reports from South Africa and the United Kingdom suggest that the severity of clinical disease with Omicron may be lower than for prior variants. However, it is unclear if this reduction in clinical severity in humans is due to the Omicron variant itself or if it is reflective of a high level of population immunity due to prior vaccination and/or infection. Syrian golden hamsters provide a robust model for severe clinical disease with SARS-CoV-2, with reproducible weight loss and pneumonia following SARS-CoV-2 infection^2-7^. To assess the pathogenicity of the SARS-CoV-2 Omicron variant, we developed two Omicron challenge stocks and assessed clinical disease, viral loads, and histopathology in hamsters.

We generated two independent SARS-CoV-2 Omicron stocks in VeroE6-TMPRSS2 cells inoculated with nasal swabs from Omicron-infected individuals. Omicron Stock 1 had a titer of 2.3×10^9^ TCID50/ml and 2.5×10^7^ PFU/ml in VeroE6-TMPRSS2 cells (EPI_ISL_7171744; Mehul Suthar, Emory University). Omicron Stock 2 had a titer of 2.3×10^8^ TCID50/ml and 3.8×10^6^ PFU/ml in VeroE6-TMPRSS2 cells (EPI_ISL_7263803; Yoshihiro Kawaoka, University of Wisconsin). Both stocks were fully sequenced.

Syrian golden hamsters (N=6/group) were inoculated by the intranasal route with 100 ul virus containing 2×10^6^ TCID50 in VeroE6-TMPRSS2 cells (5×10^4^ PFU) WA1/2020, Alpha (B.1.1.7), Beta (B.1.351), and Delta (B.1.617.2) stocks, essentially as we previously reported^2,3^. Note that TCID50 titers in VeroE6-TMPRSS2 cells are approximately 100-fold higher than previously reported TCID50 titers in VeroE6 cells. Infected hamsters showed a mean reduction of 4-10% of body weight by day 4 (**Fig. 1a**) and 10-17% of body weight by day 6 (**Fig. 1b**)^2,3^. One animal infected with WA1/2020 reached the 20% weight loss criteria for humane euthanization, but the rest of the hamsters recovered their body weights by approximately day 10.

**Figure 1.**
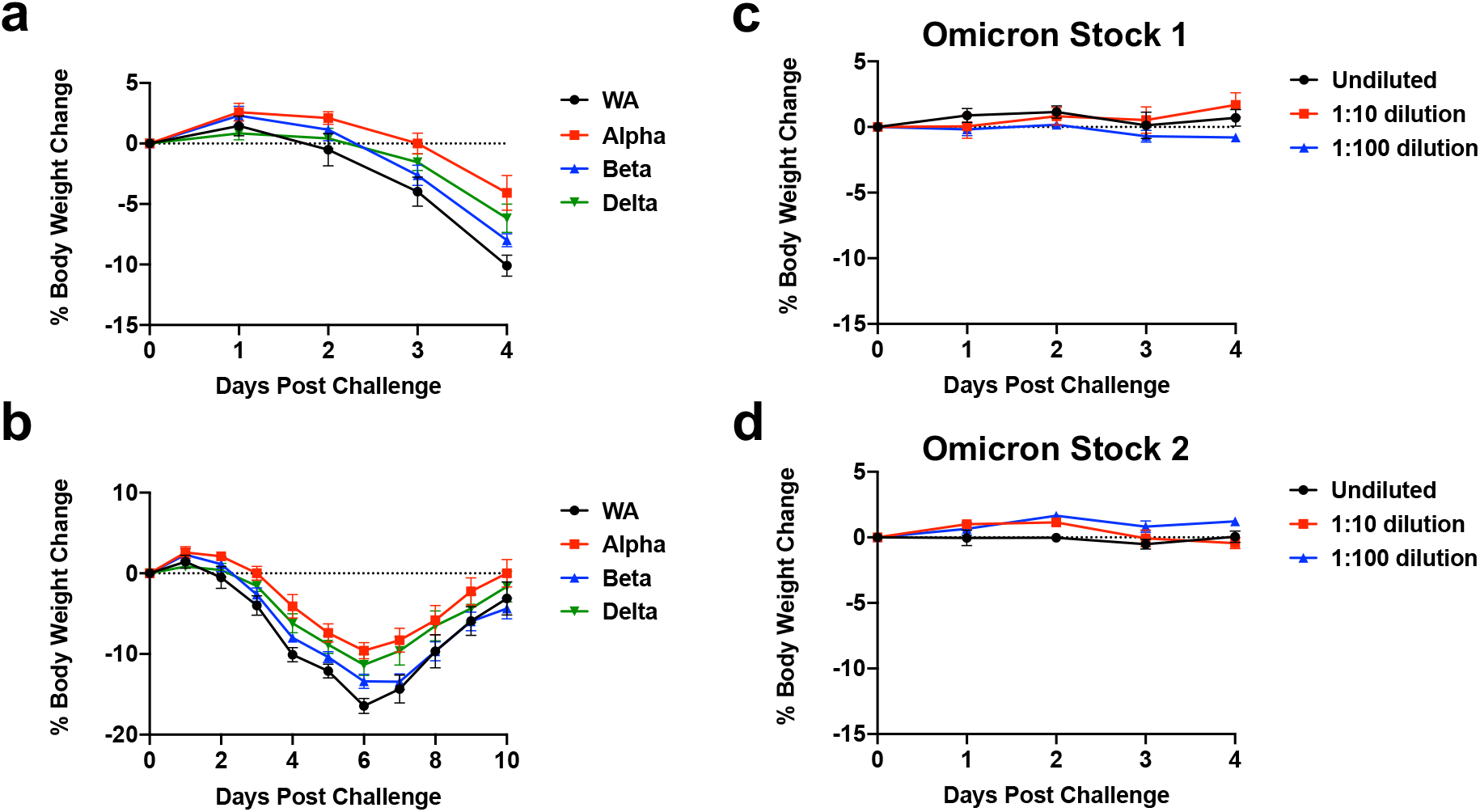
Weight loss in hamsters infected with SARS-CoV-2 variants. **a, b**, Mean body weight change following infection of hamsters with SARS-CoV-2 WA1/2020, Alpha, Beta, and Delta variants. **c, d**, Mean body weight change following infection of hamsters with SARS-CoV-2 Omicron Stock 1 and Stock 2. Mean body weight change with standard errors are shown.

Additional groups of hamsters (N=4/group) were concurrently inoculated by the intranasal route with 100 ul of undiluted, 1:10 dilution, or 1:100 dilution of Omicron Stock 1 or Stock 2. For Stock 1, the challenge doses were 2.3×10^8^ TCID50 (2.5×10^6^ PFU), 2.3×10^7^ TCID50 (2.5×10^5^ PFU), and 2.3×10^6^ TCID50 (2.5×10^4^ PFU). Hamsters inoculated with Omicron Stock 1 showed a mean reduction of -1%, -2%, and 1% of body weight by day 4 for these challenge doses, respectively (**Fig. 1c**). For Stock 2, the challenge doses were 2.3×10^7^ TCID50 (3.8×10^5^ PFU), 2.3×10^6^ TCID50 (3.8×10^4^ PFU), and 2.3×10^5^ TCID50 (3.8×10^3^ PFU). Hamsters inoculated with Omicron Stock 2 showed a mean reduction of 0%, 0%, and 1% of body weight by day 4 for these challenge doses, respectively (**Fig. 1d**). Additional follow-up time beyond day 4 did not reveal any late weight loss (data not shown). These data demonstrate that two independent SARS-CoV-2 Omicron stocks did not lead to clinical weight loss in hamsters, even when inoculated at high doses.

We next assessed tissue viral loads on day 4 following infection of hamsters with 5×10^4^ PFU WA1/2020 (N=13), 5×10^4^ PFU Beta (N=11), and the titration of Omicron Stock 1 described above (N=12). Levels of E subgenomic RNA (sgRNA) and N genomic RNA (gRNA) were assessed by RT-PCR in lungs and nasal turbinates^8-10^. In lung tissue, hamsters infected with WA1/2020, Beta, and Omicron had a median of 9.07, 9.55, and 8.33 log sgRNA copies/g tissue. Median levels of lung sgRNA were 0.74 log lower in Omicron-infected hamsters compared with WA1/2020-infected hamsters (P=0.004, two-tailed Mann-Whitney test; **Fig. 2a**). In nasal turbinates, hamsters infected with WA1/2020, Beta, and Omicron had a median of 6.35, 7.34, and 8.25 log sgRNA copies/g tissue. Median levels of nasal turbinate sgRNA were 1.90 log higher in Omicron-infected hamsters compared with WA1/2020-infected hamsters (P=0.05, two-tailed Mann-Whitney test; **Fig. 2a**). Similarly, median levels of lung gRNA were 0.84 log lower in Omicron-infected hamsters compared with WA1/2020-infected hamsters (P=0.005, two-tailed Mann-Whitney test; **Fig. 2b**), and median levels of nasal turbinate sgRNA were 1.49 log higher in Omicron-infected hamsters compared with WA1/2020-infected hamsters (P=0.01, two-tailed Mann-Whitney test; **Fig. 2b**). Tissue viral loads were similar in Omicron-infected animals at the three doses tested (2.5×10^6^, 2.5×10^5^, and 2.5×10^4^ PFU) (**Extended Data Figs. 1, 2**). These data suggest that Omicron infection in hamsters results in higher levels of virus in the upper respiratory tract but lower levels of virus in the lower respiratory tract compared with WA1/2020 infection.

**Figure 2.**
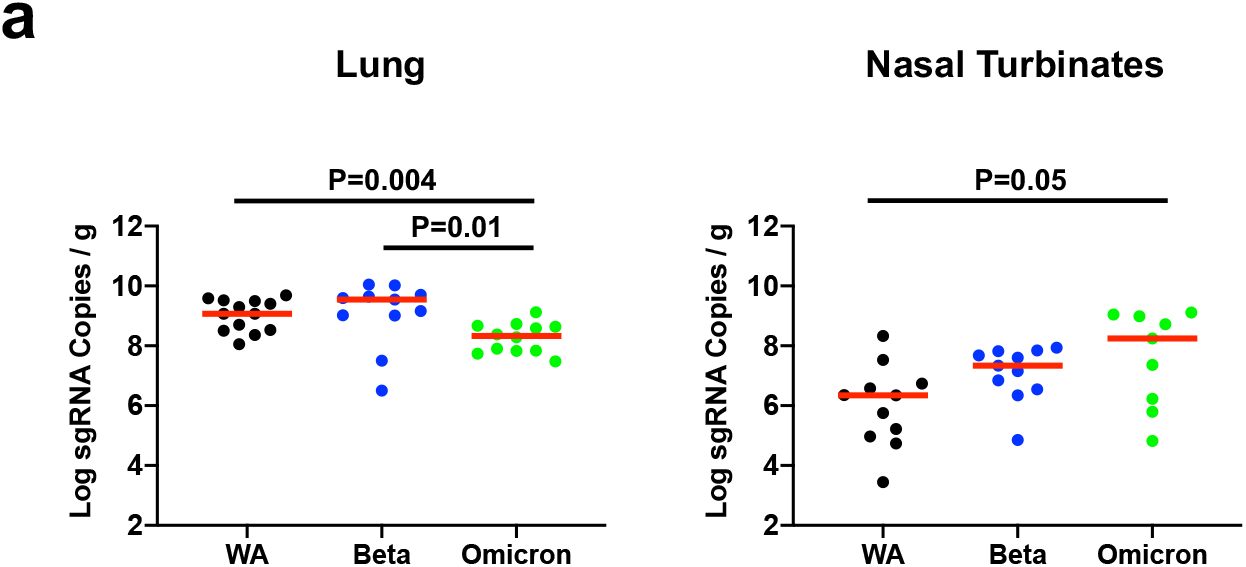
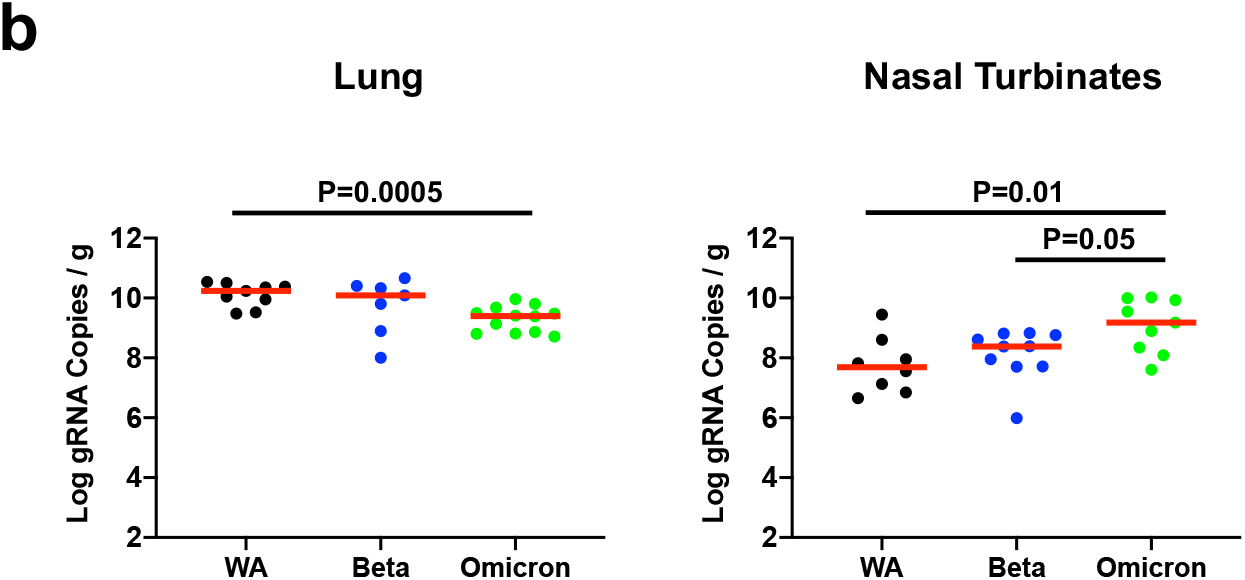
Tissue viral loads in hamsters on day 4 following SARS-CoV-2 infection. **a**, E subgenomic RNA (sgRNA) levels in lung and nasal turbinates following infection of hamsters with SARS-CoV-2 WA1/2020, Beta, and Omicron variants. **b**, N genomic RNA (gRNA) levels in lung and nasal turbinates following infection of hamsters with SARS-CoV-2 WA1/2020, Beta, and Omicron variants. Log sgRNA copies per gram tissue are shown. Medians (red bars) are depicted. P-values represent two-sided Mann-Whitney tests.

Histopathology of lung tissue on day 4 following infection with WA1/2020 or Omicron revealed multifocal areas of interstitial inflammation and consolidation associated with bronchioles (**Fig. 3a**). Bronchiolar epithelium showed degeneration, loss of cilia, loss of nuclear polarity, and sloughing in both WA1/2020 and Omicron infected hamsters (**Fig. 3b**). Vascular inflammatory infiltrates and endothelialitis were observed in both WA1/2020 and Omicron infected hamsters (**Extended Data Fig. 3**).

**Figure 3.**
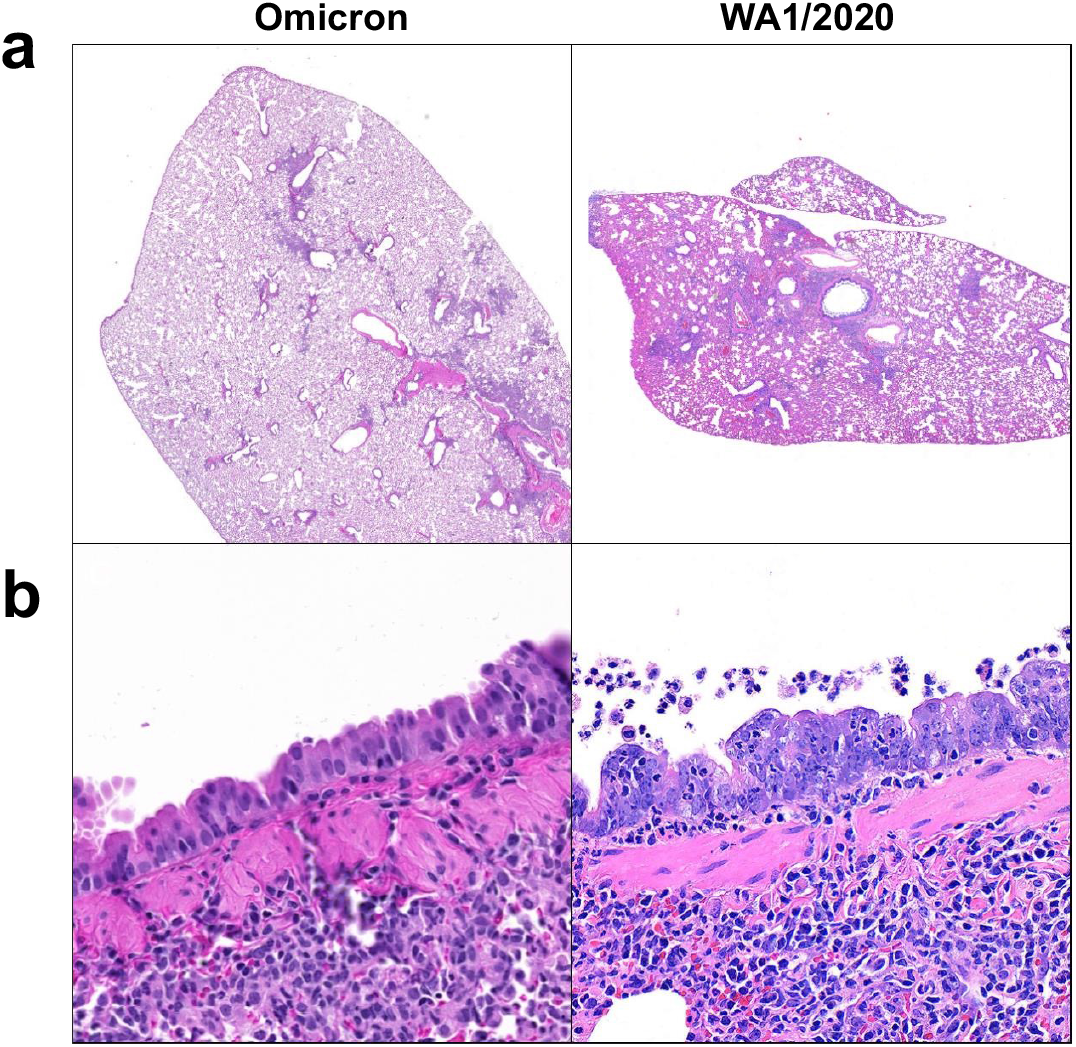
Histopathology in hamsters on day 4 following SARS-CoV-2 infection. Lung tissue from hamsters infected with SARS-CoV-2 WA1/2020 and Omicron variants was stained with H&E. **a**, Low power representative images of lung. There are multifocal areas of interstitial inflammation and patchy consolidation associated with bronchioles. **b**, Medium power representative images of bronchiolar epithelium. There is degeneration of the epithelium characterized by loss of cilia, loss of nuclear polarity, and sloughing into the airway, with consolidation in the subjacent pulmonary parenchyma.

Our data demonstrate that the SARS-CoV-2 Omicron variant infects Syrian golden hamsters over a broad range of challenge doses. However, Omicron infection did not result in detectable weight loss, even at doses that were 100-times higher than the doses of WA1/2020, Alpha, Beta, and Delta that led to substantial weight loss. The consistency of these findings with two independent Omicron challenge stocks suggests the generalizability of these conclusions. Omicron infection led to robust viral replication in both the upper and lower respiratory tracts, but with higher viral loads in nasal turbinates and lower viral loads in lung compared with WA1/2020. These data suggest that Omicron infection may lead to increased upper respiratory tract disease but reduced lower respiratory tract disease. Nevertheless, Omicron infection still led to lung pathology, including patchy consolidation, bronchiolar epithelial degeneration, and endothelialitis.

Our findings in hamsters are consistent with preliminary reports suggesting that Omicron is more transmissible but induces less severe clinical pneumonia in humans compared with prior SARS-CoV-2 variants. If confirmed, these data would have major public health implications given the trajectory of the global Omicron surge. Future studies could further define the pathogenesis of infection with the SARS-CoV-2 Omicron variant in hamsters and the molecular mechanisms associated with its reduced pathogenicity and apparent skewing towards increased upper respiratory tract disease and decreased lower respiratory tract disease.

## Acknowledgements

The authors acknowledge NIH grant CA260476, the Massachusetts Consortium for Pathogen Readiness, the Ragon Institute, and the Musk Foundation (D.H.B.).

## Data sharing

All data are available in the manuscript or the supplementary material. Correspondence and requests for materials should be addressed to D.H.B.

## Author Contributions

This study was designed by DHB. Omicron challenge stocks were prepared by MS and YK. Animal work was performed by HA and MGL. Virologic assays were performed and analyzed by KM, VG, LHT, BC, and MS. Histopathology was performed by CPM and AJM.

## Methods

### Study design

Eight week old female and male golden Syrian hamsters (Envigo) were infected with SARS-CoV-2 in a volume of 100 µl (50 µl/nostril) by the intranasal route. Following challenge, body weights were assessed daily. Body weight loss that exceeded 20% of the weight on the day of challenge was established as a humane endpoint euthanasia criteria. A subset of animals were necropsied on day 4 for tissue viral loads and histopathology. All animal studies were conducted in compliance with all relevant local, state, and federal regulations and were approved by the BIOQUAL Institutional Animal Care and Use Committee.

### SARS-CoV-2 viral stocks

The WA1/2020 (USA-WA1/2020; BEI Resources NR-5228), Alpha (B.1.1.7; USA/CA_CDC_5574/2020; BEI Resources NR-54011), Beta (B.1.351; South Africa/KRISP-K005325/2020; BEI Resources NR-54974), and Delta (B.1.617.2; USA/PHC658/2021; BEI Resources NR-55612) challenge stocks have been previously described^2^. We generated two independent SARS-CoV-2 Omicron stocks in VeroE6-TMPRSS2 cells inoculated with nasal swabs from Omicron-infected individuals. Omicron Stock 1 had a titer of 2.3×10^9^ TCID50/ml and 2.5×10^7^ PFU/ml in VeroE6-TMPRSS2 cells (EPI_ISL_7171744; Mehul Suthar, Emory University). Omicron Stock 2 had a titer of 2.3×10^8^ TCID50/ml and 3.8×10^6^ PFU/ml in VeroE6-TMPRSS2 cells (EPI_ISL_7263803; Yoshihiro Kawaoka, University of Wisconsin). All SARS-CoV-2 stocks were fully sequenced. All TCID50 and PFU assays were in VeroE6-TMPRSS2 cells.

### Genomic and subgenomic viral load assays

SARS-CoV-2 N gene genomic RNA (gRNA) and E gene subgenomic RNA (sgRNA) was assessed by reverse transcription polymerase chain reactions (RT-PCR) using primers and probes as previously described.^8-10^ Standards were generated by first synthesizing a gene fragment of the genomic N gene or subgenomic E gene.^9^ The gene fragments were subsequently cloned into a pcDNA3.1+ expression plasmid using restriction site cloning (Integrated DNA Technologies). The inserts were in vitro transcribed to RNA using the AmpliCap-Max T7 High Yield Message Maker Kit (CellScript). Log dilutions of the standard were prepared for RT-PCR assays ranging from 1×10^10^ copies to 1×10^-1^ copies. Viral loads were quantified from lung tissue as follows; total RNA was extracted on a QIAcube HT instrument using the RNeasy 96 QIAcube HT Kit according to manufacturer’s specifications (Qiagen). Standard dilutions and extracted total RNA from samples were reverse transcribed using SuperScript VILO Master Mix (Invitrogen) according to manufacturer’s specifications. A Taqman custom gene expression assay (Thermo Fisher Scientific) was designed using the sequences targeting the E gene sgRNA.^9^ The sequences for the custom assay were as follows, forward primer, sgLeadCoV2.Fwd: CGATCTCTTGTAGATCTGTTCTC, E_Sarbeco_R: ATATTGCAGCAGTACGCACACA, E_Sarbeco_P1 (probe): VIC-ACACTAGCCATCCTTACTGCGCTTCG-MGB. For the genomic N assays, the sequences for the forward (F) and reverse (R) primes and probe (P) were: 2019-nCoV_N1-F :5’-GACCCCAAAATCAGCGAAAT-3’; 2019-nCoV_N1-R: 5’-TCTGGTTACTGCCAGTTGAATCTG-3’; 2019-nCoV_N1-P: 5’-FAM-ACCCCGCATTACGTTTGGTGGACC-BHQ1-3’. Reactions were carried out in duplicate for samples and standards on the QuantStudio 6 and 7 Flex Real-Time PCR Systems (Applied Biosystems). The following thermal cycling conditions were used; initial denaturation at 95°C for 20 seconds, then 45 cycles of 95°C for 1 second and 60°C for 20 seconds. Standard curves were used to calculate subgenomic RNA copies and copy number was normalized to the input weight of lung tissue (copies/g); the quantitative assay sensitivity was 100 copies per gram.

### Histopathology

Tissues were fixed in freshly prepared 4% paraformaldehyde for 24 hours, transferred to 70% ethanol, paraffin embedded within 7 to 10 days, and block sectioned at 5 µm. Slides were baked for 30 to 60 min at 65 °C and then deparaffinized in xylene and rehydrated through a series of graded ethanol to distilled water. Slides were stained with hematoxylin and eosin. Blinded assessment of tissue pathology was performed by a veterinary pathologist (AJM).

### Statistical analyses

Analysis of virologic and body weight data was performed using GraphPad Prism 9.1.2 (GraphPad Software). Comparison of data between groups was performed using two-sided Mann–Whitney tests. *P* values of less than 0.05 were considered significant.

## Extended Data Figure Legends

**Extended Data Figure 1.**
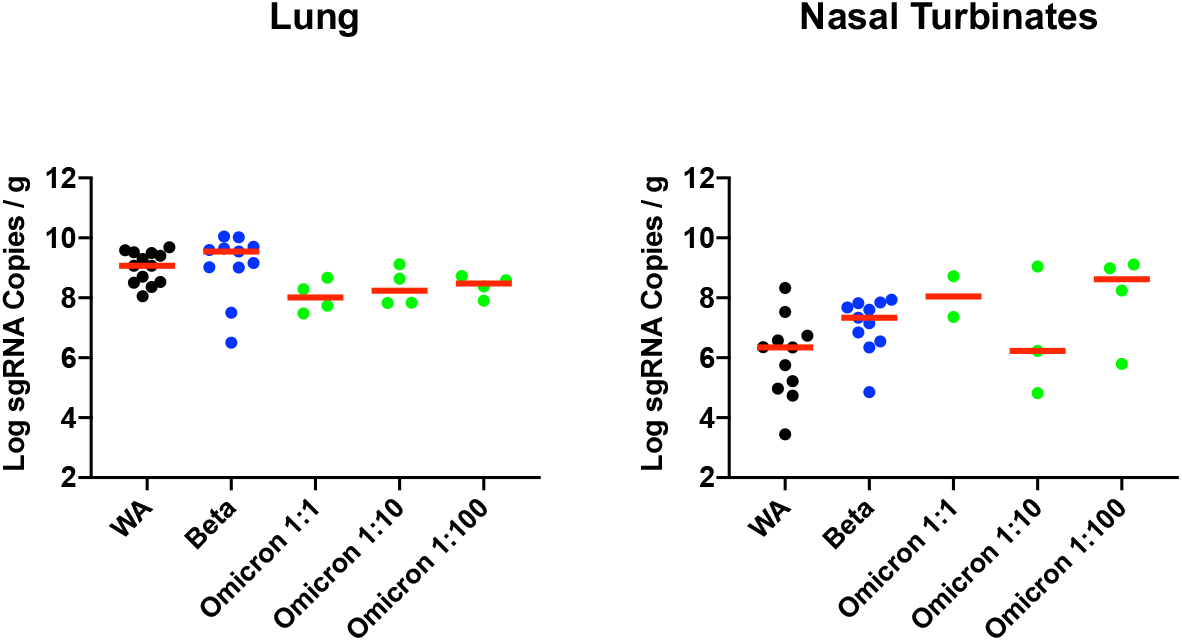
Tissue sgRNA in hamsters on day 4 following SARS-CoV-2 infection. E subgenomic RNA (sgRNA) levels in lung and nasal turbinates following infection of hamsters with SARS-CoV-2 WA1/2020, Beta, and a titration of doses of Omicron. Log sgRNA copies per gram tissue are shown. Medians (red bars) are depicted.

**Extended Data Figure 2.**
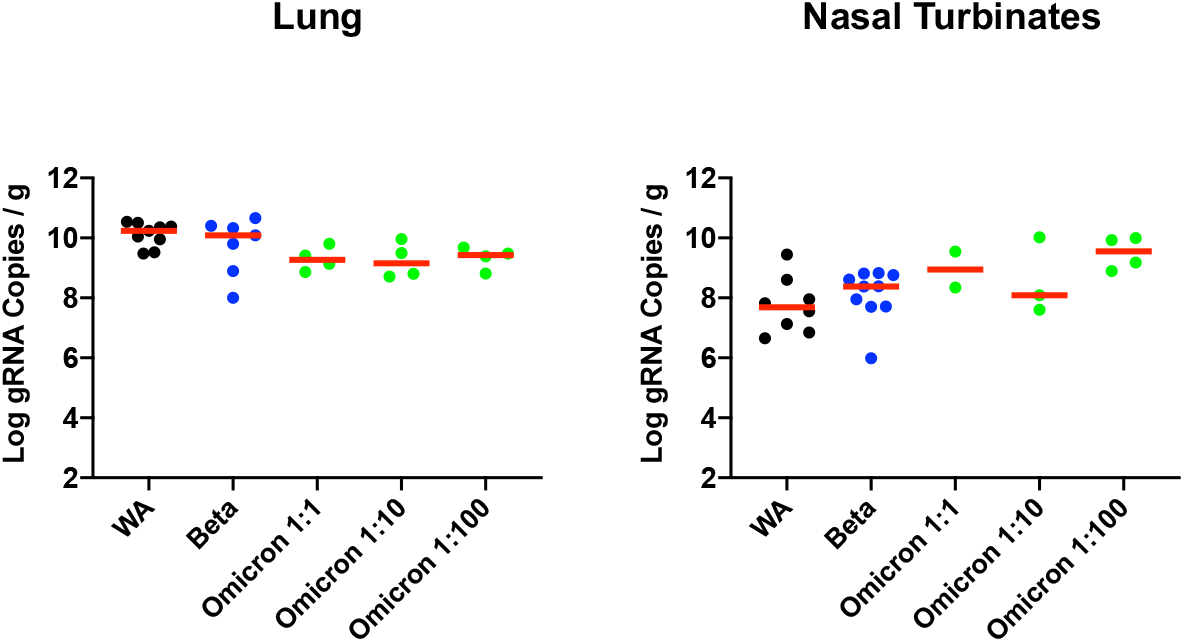
Tissue gRNA in hamsters on day 4 following SARS-CoV-2 infection. N genomic RNA (gRNA) levels in lung and nasal turbinates following infection of hamsters with SARS-CoV-2 WA1/2020, Beta, and a titration of doses of Omicron. Log gRNA copies per gram tissue are shown. Medians (red bars) are depicted.

**Extended Data Figure 3.**
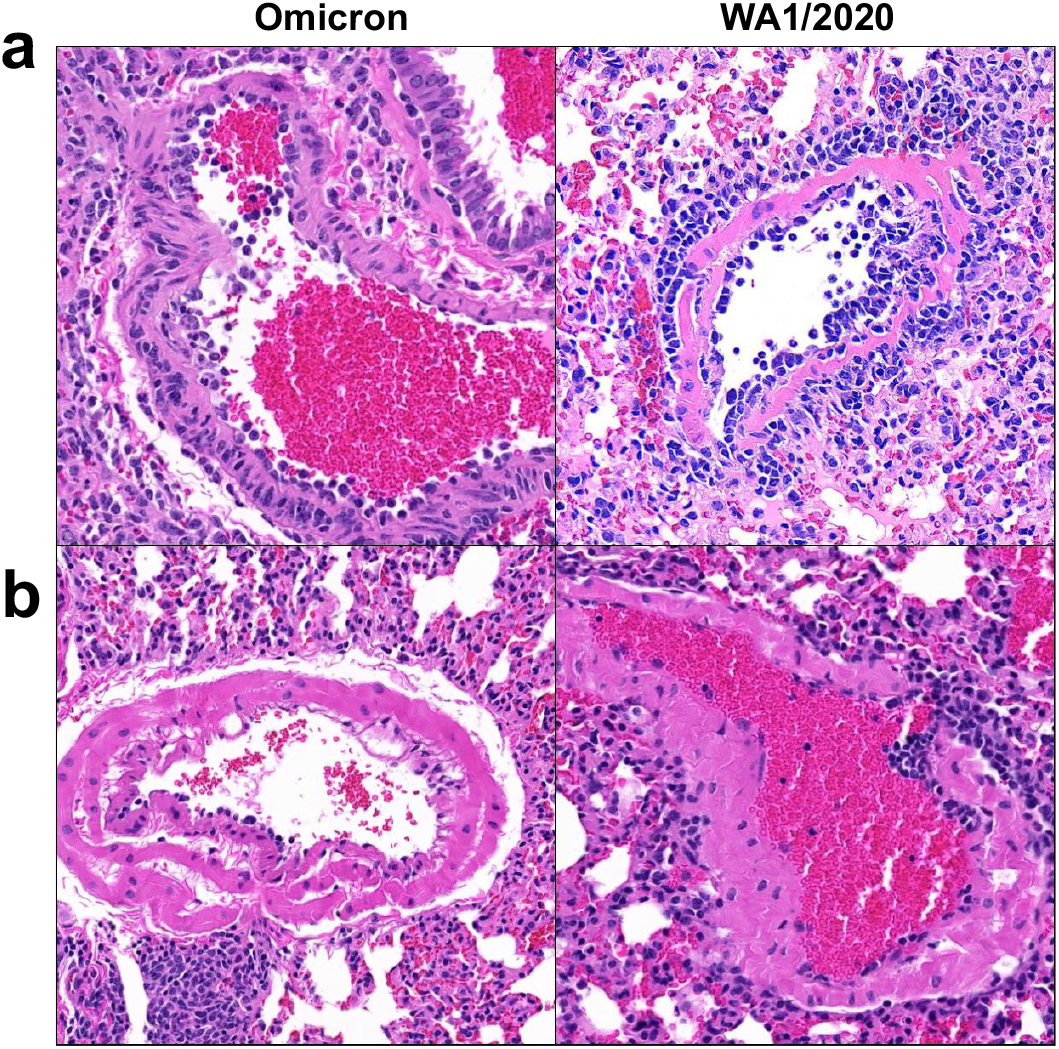
Vasculitis in hamsters on day 4 following SARS-CoV-2 infection. Lung tissue from hamsters infected with SARS-CoV-2 WA1/2020 and Omicron variants was stained with H&E. **a**, Pulmonary arteries showing vascular inflammatory infiltrates and endothelialitis. **b**, Similar but decreased pathology observed at the 1:100 dilution challenge dose of the Omicron variant.

